# Ongoing local adaptation to climate change in yarrow (*Achillea millefolium* L.)

**DOI:** 10.1101/2024.07.22.604595

**Authors:** Gianalberto Losapio, Baptiste Doussot, Fabrizio Araniti, Leonardo Bruno, Roger Guevara, Rodolfo Dirzo

**Affiliations:** Department of Biology, Stanford University, USA; Institute of Earth Surface Dynamics, University of Lausanne, Switzerland; Department of Biosciences, University of Milan, Italy; Department of Agricultural and Environmental Sciences, University of Milan, Italy; Department of Biology, Ecology and Earth Sciences, University of Calabria, Italy; Red Biología Evolutiva, Instituto de Ecologia, AC, Mexico; Department of Earth Systems Science, Stanford University, USA

## Abstract

Climate change threatens biodiversity as populations can persist if they migrate or adapt to the rapidly changing conditions of the Anthropocene. However, little is known about the persistence of plant populations under the long-term trends of increasing temperature and drought. We explore whether historical populations of yarrow (*Achillea millefolium* L.) surveyed in the 1920s have persisted locally or undergone local extinction after 100 years. We resurveyed historical sites spanning a broad climatic gradient (from 1 m to 3,200 m a.s.l.), examined to what extent plant growth signature (height) has changed over time, and analyzed metabolic diversity. Nine out of ten populations persisted locally and showed phenotypic and metabolic differentiation. The only population potentially extirpated is that of the hottest and driest site. Populations from warm sites in coastal and mountain regions had grown taller than 100 years ago. In contrast, populations from dry sites in lowlands and foothills became shorter. Furthermore, we document differentiation in metabolic diversity involving plant defenses and signaling. Ongoing local adaptation is constrained by changing both temperature and precipitation as well as by biotic interactions. Preserving locally adapted populations and metabolic diversity is key for conservation efforts.

## 1. Introduction

Climate change is a complex alteration of global and local climatic patterns compared to a last century-based line which includes warming and precipitation regime shifts (IPCC 2022). Considerable effort has been invested in understanding the effects of rising temperatures on organisms and populations (Parmesan & Hanley, 2015; Steinbauer et al., 2018; Zandalinas et al., 2021; Manrique-Ascencio et al., 2024). Yet, besides temperature, the long-term effects of increasing drought remain unclear (Losapio & Schöb, 2017; Exposito-Alonso et al., 2018). As the acceleration of these changes in climate poses new challenges to species’ survival and conservation (Parmesan & Hanley, 2015), understanding species’ response to rapid environmental change is urgent and critical to guide policy for halting the biodiversity loss crisis (Dirzo et al. 2022).

The seminal, pioneering work of Hall, Clausen, Keck, and Hiesey (Clausen et al., 1948) documented for the first time the adaptive responses of species to changing environments. Through field observations and common-garden experiments, Clausen et al. (1948) demonstrated that geographically distinct populations of yarrow (*Achillea millefolium* L.) have adapted to specific local climate conditions. They documented how certain traits, particularly plant height and phenology, vary across geographical gradients in response to changes in temperature and precipitation (Clausen et al.1948). They proposed that yarrow ecotypes possess varying degrees of genetic diversity, which contributes to their local adaptation and broad distribution across different environments. Today, after one hundred years of increasing temperature and drought, it remains unknown if those locally adapted populations persisted or went extinct and whether the growth signature (plant height) has changed in response to locally changing climatic conditions.

As climate change accelerates, it poses new challenges to species’ survival and conservation (Parmesan & Hanley, 2015). Genetic diversity is key to the ability of populations to survive, reproduce, and adapt under changing climatic conditions (Kawecki & Ebert, 2004; Liu et al., 2019; Bastias et al., 2024). Associated with genetic diversity, chemical diversity plays a central role in species’ potential to adapt to changing climatic conditions and biotic environments (Agrawal et al., 2009; Wetze and Whitehead, 2020; Walker et al., 2023). Also, plant secondary metabolism may be affected by changes in climatic factors (Fernandez-Conradi et al., 2022). Specialized plant metabolites have key functions in mediating species interactions and driving the response of plants to environmental stress (Agrawal et al., 2009; Fernandez-Conradi et al., 2022; Volf et al., 2024). However, how the potential contribution of chemical diversity to local adaptation changes over space-time remains poorly documented. Understanding the metabolic mechanisms underlying local adaptation is essential for anticipating species responses to climate change (Agrawal et al., 2009; Walker et al., 2023). This knowledge is crucial for devising effective conservation and restoration strategies that safeguard biodiversity and ecosystem integrity.

In this article, we explore whether historical populations of yarrow persist or not after one hundred years, and if/how they have phenotypically changed. We resurveyed ten locally-adapted yarrow populations that were originally surveyed in the 1920s (Clausen et al., 1948) to compare them with current populations from the same locations. We then established a common garden experiment to assess the metabolic diversity of different populations under the same controlled climatic conditions. By means of metabolomics, we examined if populations from different regions exhibit inherited metabolic adaptations that may have allowed them to persist over one hundred years of increasing temperature and drought. We address the following questions: (1) Do yarrow populations persist or are they locally extirpated? (2) Have yarrow populations changed regarding plant height (signature trait) over the last 100 years? (3) Which metabolic components predict differences among locally-adapted populations?

## 2. Material and methods

### (a) Re-survey data

To address the first question on population persistence, we revisited historical study sites of yarrow local-adaptation studies reported by Clausen et al. (1948) during the spring and summer of 2020. We selected the ten sites where experimental evidence of local adaptation was documented (electronic supplementary information, table S1). These ten sites span a broad climatic gradient in California, ranging from the Pacific coast at 1 m a.s.l. to the alpine zone at 3,200 m a.s.l. In each site, we (1) recorded the occurrence of the yarrow population, (2) measured plant height (1 cm accuracy from bottom to top of inflorescence) on ten randomly chosen plants in reproductive phase, and (3) collected seeds.

### (b) Data analysis

To address the second question on population changes over space-time, we extracted the original plant height data for each site as reported from the 1920s studies (Clausen et al. 1948; electronic supplementary information, table S2). We re-sampled plant height data from two uniform distributions. Twenty plants were randomly sampled from the min–max interval of each site for the two years (i.e., the 1920s and 2020). We gathered climate data for both 1920 and 2020 for mean annual temperature and annual precipitation from https://calclim.dri.edu/

We used a linear regression model to test for differences in plant height (response variable) between years (categorical predictor), temperature, precipitation, and their statistical interactions. To this end, we used the *lm* function of base R software package (R Core team 2023). The model was evaluated in terms of residuals dispersion, variance explained, and Cohen’s effect size was calculated (electronic supplementary information, R code) (R Core team 2023).

We first tested for differences in plant height among sites between years using a linear regression model. We estimated marginal (least-squares) means for each combination of site and year (*n* = 18) and compared plant height within each site between years. We used the *emmeans* function of the R software package of the same name (Lenth 2023). Then, we analyzed the possible change of plant height over time using a stochastic ecological model of plant growth in response to climate change (Soetaert and Herman, 2009), which was parametrized using field survey data. Changes in plant height *H* of each population *i* over time *t* were modeled as

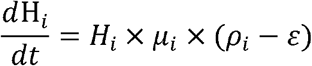

where *µ* represents plant growth rate and *p* the plant response to climate between 1920 and 2020; the stochastic component was represented by *ε* as random mortality (for more details, see electronic supplementary material, supplementary methods b).

### (c) Metabolomic analysis

To address the third question on plant metabolic diversity among populations, we sowed 1 g of seeds per ecotype in germination trays with well-watered potting soil in an open-chamber greenhouse. Germination rates were recorded every day for the first two weeks and then once per week. After 60 days, we transplanted one plant per pot (common potting soil in 8 cm pots) and let the yarrow grow for another 60 days before metabolomic analysis. We analyzed the ecotypes of Bodega Bay, Groveland, Mather, and Yosemite Creek, representing the Pacific coast, low, medium, and high elevation adaptation, respectively.

Plant secondary metabolites, particularly volatile organic compounds (VOCs), were analyzed via Headspace Solid-Phase Microextraction coupled to Gas Chromatography-Mass Spectrometry (HS-SPME-GC-MS). We used an Agilent gas chromatograph (GC 7890A) coupled with a single quadrupole mass spectrometer (MS 5975C INERT XL MSD) and a CTC ANALYTICS PAL autosampler. To perform sample chromatography, a 5MS column (30 m×0.25 mm×0.25µm + 10 m of pre-column) was employed (electronic supplementary material, supplementary methods a). For the experiments, one gram of fresh plant material was used. Before injection, the samples were equilibrated for 20 minutes, at room temperature, in a 20 ml sealed glass vial.

Metabolomic analysis was performed using MetaboAnalyst 5.0 (Pang *et al*., 2022). Metabolomic data were first normalized using the MS-DIAL mTIC (Total Ion Current). We used Partial Least-Squares Discriminant Analysis (PLS-DA) to distinguish populations and identify compounds most contributing to those differences. Then, we used a machine learning algorithm (i.e., Random Forest) to select compounds with the highest discriminatory power; a threshold > 1 was set for variable importance in the projection (VIP) score (Pang *et al*., 2022). The PLS-DA model was validated using Q2 as a performance measure through 10-fold cross-validation. Chemical information was retrieved from open chemistry database at the National Institutes of Health (Kim et al., 2023).

## 3. Results

Climatic conditions have changed over the last one hundred years in all sites (Table S1, electronic supplementary information). The temperature increased monotonically across sites, with changes between +4.63% and +23.82%; changes were more pronounced at high-elevation sites than at lower-elevation. On the contrary, changes in precipitation were idiosyncratic as precipitation increased in coast sites while decreased in lowlands and western slopes of the Sierra Nevada (Table S1, electronic supplementary information).

Yarrow populations occurred at nine out of ten historical sites. The only population not recorded, potentially extirpated, was that of the hottest and driest site in the lowland (Knights Ferry). Inspection of this site did not reveal habitat conversion due to land use change or any other visible disturbance (personal observation).

Overall, yarrow plants grew taller over one hundred years on average across sites (coef: 1.04 ± 0.25, p < 0.001; electronic supplementary information). Changes in yarrow plant height were significantly correlated to changes in temperature (coef: 0.09 ± 0.02, p < 0.001; fig. 1a), precipitation (coef_linear_: -22.89 ± 4.29, p < 0.001; coef_quadratic_: 9.57 ± 5.35, *p* = 0.074; fig. 1b), and by the interaction temperature–precipitation–years (coef_linear_: -1.32 ± 0.42, p = 0.002; coef_quadratic_: -2.40 ± 0.50, p < 0.001).

**Figure 1.**
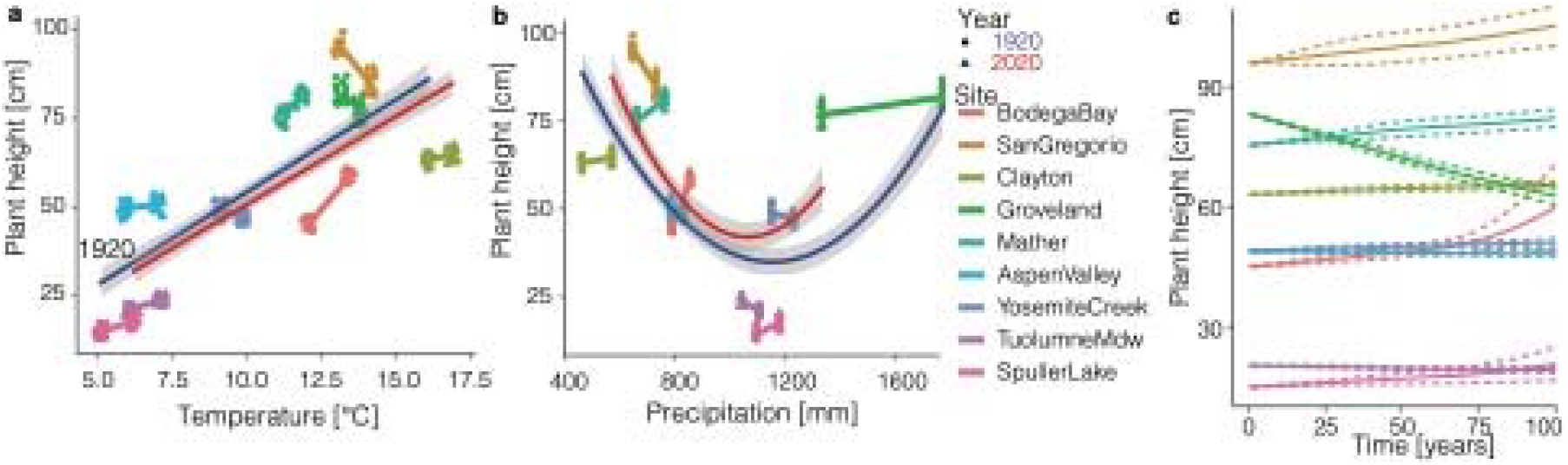
Changes in yarrow plant height (*y*-axis) in response to changes in temperature (a) and precipitation (b) between the 1920s (blue line and circle points) and 2020 (red line and triangle points). The line connecting the two average data points indicate the strength and direction of the change between years. (c) A model of plant height evolution over one hundred years of changing temperature and precipitation across populations.

The degree of change in yarrow height between years varies among populations (f_years:site_ = 1.28, p < 0.001; electronic supplementary information). Between 1920s and 2020, yarrow plants grew significantly taller at Bodega Bay (contrast: 0.261 ± 0.015, p < 0.001), Mather (contrast: 0.076 ± 0.015, *p* < 0.001), Aspen Valley (contrast: 0.029 ± 0.015, p = 0.047), Tuolumne Meadow (contrast: 0.125 ± 0.015, p < 0.001), and Spuller Lake (contrast: 0.193 ± 0.015, p < 0.001). On the contrary, yarrow plants were significantly shorter at San Gregorio (contrast: -0.121 ± 0.015, p < 0.001), Groveland (contrast: -0.063 ± 0.015, p < 0.001), and Yosemite Creek (contrast: 0.039 ± 0.015, p = 0.009). Yarrow plants were not statistically different between years at Clayton (contrast: 0.023 ± 0.015, p = 0.121).

Results of a theoretical model parametrized with field data indicate, in seven out of nine cases, that changes in yarrow height are driven by local adaptation to changes in both temperature and precipitation (f_years:site_ = 0.49, *p* < 0.001; fig. 1c).

The metabolomic analysis identified 102 plant compounds, 42 of which were found in significantly different concentrations among populations (p < 0.001). Overall, yarrow populations showed distinct metabolic profiles (Fig. 2a). Random forest results indicate 14 metabolites predict differences in metabolic profile (all VIP scores > 1.6). In particular, the concentrations of 4-Terpinyl acetate, Isobornyl formate, p-Menth-3-en-1, Dehydrosabinenol, cis-3-Hexenyl, and α-Terpinen were the highest in high-elevation populations (Fig. 2b). In contrast, the concentrations of Santolina triene, Artemisia ketone, 3-Methyl-4-pentenal, and Myrtenyl acetate were the highest in low-elevation populations. The concentration of Camphor, p-Mentha-1,4-diene, 2,3-Dehydro-1,8-cineole and Eucalyptol was the highest in populations from mid-elevation.

**Figure 2.**
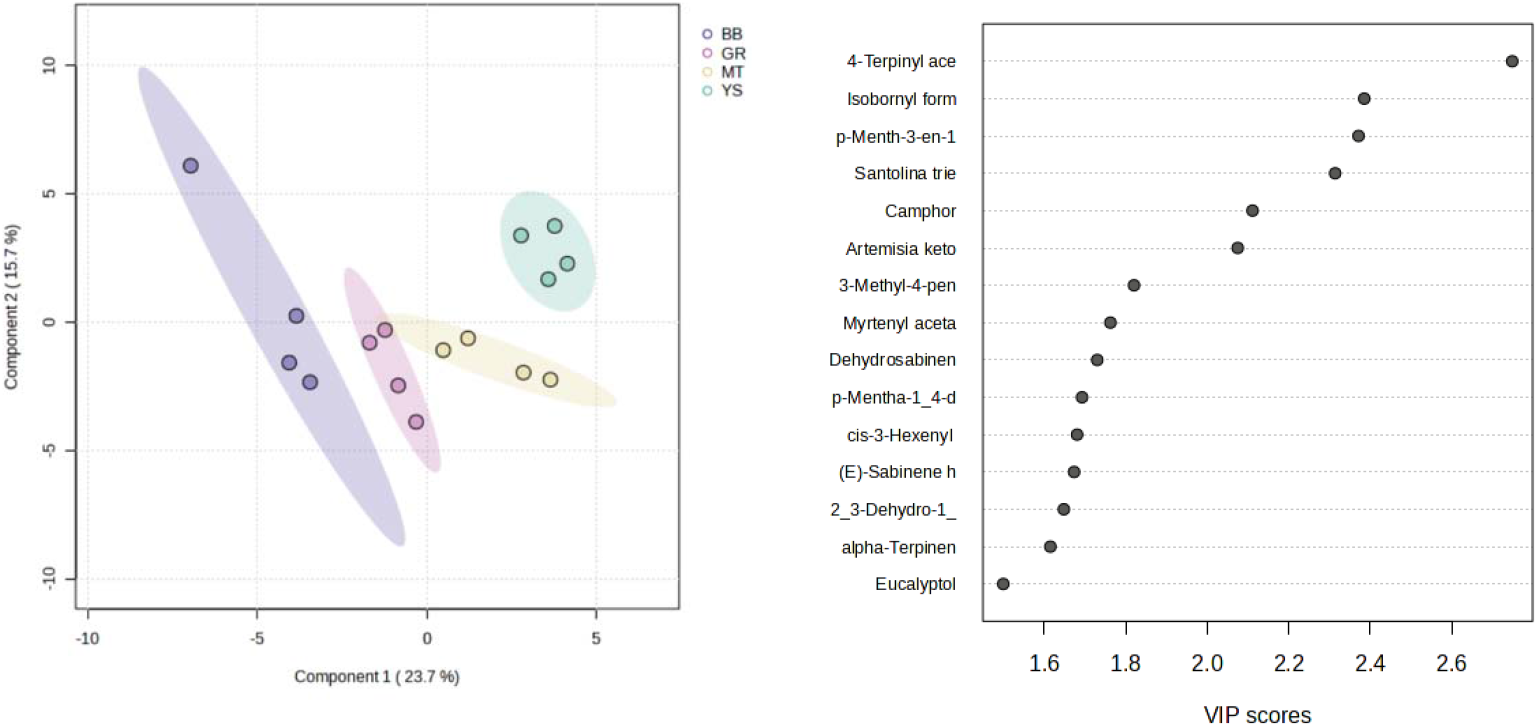
Biplot of PLS-DA reporting the distribution of populations (BB = Bodega Bay; GR = Groveland; MT = Mather; YS = Yosemite Creek) along the first two components. Panel on the right shows the most important metabolites responsible for differentiating populations.

## 4. Discussion

Our results provide novel insights into the responses of yarrow populations (*Achillea millefolium* L.) to one hundred years of climate change. We report that nine out of ten historical populations were still viable, indicating the ongoing local adaptation of this species. However, the population of Knights Ferry, the hottest and driest site, may have been potentially extirpated, highlighting the vulnerability of populations to increasing both temperature and aridity at the range edge (Bastias et al., 2024). There, extirpation might have been driven by additional factors other than climate overlooked here.

We document an increase in plant height over one hundred years in most populations parallel to a warming climate. Our results suggest that yarrow plants tend to grow taller as temperature has increased during the last century. This finding is consistent with global trends in the growth–temperature relationship, whereby plants become taller with increasing temperature or, vice versa, become shorter with increasing altitude or latitude (Buckley & Puy, 2022).

Besides rising temperatures, many populations also experienced a dramatic change in precipitation regimes. Consistently, we found a complex, non-linear relationship between yarrow growth and precipitation patterns. Significant interactions between temperature and precipitation were found as the effects of increasing temperature depend on how precipitation changes: as temperature and drought increased, yarrow plants became shorter. An ecological model simulation indicates that yarrow plants get shorter if precipitation decreases while temperature increases. These results are consistent with physiological constraints, as increasing temperature at constant precipitation leads to higher evapotranspiration and, hence water deficit (Parmesan & Hanley, 2015)

Identifying distinct phytochemical profiles among populations provides further evidence supporting ongoing local adaptation and differentiation. We identified diverse sets of plant secondary metabolites that play crucial roles in ecological interactions and responses to environmental stress (Agrawal et al., 2009; Fang et al., 2019; Walker et al., 2023). The phytochemical 4-Terpinyl acetate may contribute to defense against herbivores and fungal pathogens (Kim et al., 2023). Isobornyl formate may be involved in the yarrow’s response to abiotic stressors by acting as a signaling molecule (Kim et al., 2023). p-Menth-3-en-1 contributes to the yarrow aromatic scent (Kim et al., 2023). Dehydrosabinenol can have allelopathic effects impacting the growth and survival of neighbouring plants (Kim et al., 2023). cis-3-Hexenyl is a volatile terpenoid that might be involved in response to herbivory, attracting natural enemies of herbivores and communicating with neighbouring plants (Kim et al., 2023). alpha-Terpinen has microbial properties. Santolina triene contributes to drought and high-temperature resistance (Kim et al., 2023). Artemisia ketone and Myrtenyl are volatile biomarkers acting as signaling molecules; Methyl-4-pentenal may have allelopathic effects (Kim et al., 2023). Camphor has insecticidal properties, and Eucalyptol is an antimicrobial (Kim et al., 2023); p-Mentha-1_4-diene and 2,3-Dehydro-1,8-cineole might contribute to plant signaling and communication (Kim et al., 2023).

These compounds contribute to plant defenses against herbivores and pathogens. They may also play a role in competitive interactions with other plant species and mutualistic interactions with pollinators, microbes and beneficial insects. As these compounds are often specific, our results suggest that yarrows may adapt and differentiate to changes in local biotic communities too, in addition to climate. Future studies shall address how species interactions vary across yarrow populations. Environmental conditions could also strongly impact the production of specialized metabolites involved in plant protection. Among them, plant VOCs play crucial roles in plant adaptation to the environment and act as infochemicals in interactions involving multiple trophic levels (Agrawal et al., 2009; Walker et al., 2023). The production of these compounds, which is pivotal for plant adaptation, may be influenced by global climate change factors like elevated atmospheric carbon dioxide, ozone, and temperature. This critical aspect of plant responses to climate change warrants further research.

The documented differences in plant height and metabolic profile among yarrow populations highlight how these populations may have evolved environmental-specific responses leading to their persistence over one hundred years of increasing temperature and drought. This adaptive variation in growth and metabolism may drive population differentiation and diversification (Kawecki & Ebert, 2004; Liu et al., 2019), a process essential for the distribution and persistence of populations across diverse changing climatic conditions (Parmesan & Hanley, 2015). Understanding these drivers and the consequences of local adaptation has broader implications for conservation biology. Recognizing plant populations’ metabolic diversity traits is key to making informed decisions regarding ecological restoration projects. Utilizing locally adapted populations can increase restoration efforts’ success and enhance restored ecosystems’ resilience in the face of climate change.

## Ethics

Scientific research and collection permits were obtained from United States Department of Interior, National Park Service (permit number YOSE-2020-SCI-0055).

## Data accessibility

Data and R software code will be made publicly and freely available at Zenodo upon acceptance.

## Supplemental information

Supporting information are available as electronic supplementary material.

## Declaration of AI use

We have not used AI-assisted technologies in creating this article.

## Acknowledgments

GL was financially supported by the Swiss National Science Foundation (PZ00P3_202127) and acknowledges the financial support of the European Union – NextGenerationEU, Italian Ministry of University and Research (PRIN 2022 PNRR P2022N5KYJ). GL acknowledges the technical help of Lilian Dutoit and the emotional support from Mauro Losapio. We thank Yosemite National Park Service for issuing sampling permits. We thank Marco Caccianiga, Mario Beretta, Valerio Parravicini, and Diego De Nisi for facilitating and supporting our research at University of Milan, Città Studi Botanical Garden.

## Author contributions

GL, RG, and RD conceived the study. GL conducted field work. BD conducted lab work with help from FA and LB. GL analyzed the data with input from BD. GL drafted the manuscript. All authors commented and edited the manuscript.

## Conflict of interest declaration

We declare we have no competing financial interests.

## RESOURCE AVAILABILITY

## Lead Contact

Further information and requests should be directed to and will be fulfilled by the Lead Contact, Gianalberto Losapio.

## Materials Availability

All specimens are stored in the institutional collections (Città Studi Botanic Garden of the University of Milan) and will be made available by the Lead Contact upon reasonable request.

## Notes

### Competing Interest Statement

The authors have declared no competing interest.

