## Supplemental Information for "Ongoing local adaptation to climate change in yarrow (*Achillea millefolium* L.)"

**Supplementary Methods**

a. **Phytochemical analysis**

VOCs sampling was accomplished by exposing a grey SPME fiber (StableFlex, divinylbenzene/Carboxen on polydimethylsiloxane coating; 50/30 μm coating; Supleco) to the headspace of the vial for 20 minutes at room temperature before injecting it into the GC-MS apparatus. Sample separation was achieved using the following temperature ramp: starting at 50°C, the temperature increased by 3°C per minute until reaching 200°C, followed by an isocratic hold at 200°C for 3 minutes. To calculate the retention time index (RI), an alkane mixture (C8-C30) was injected at the predetermined time.

The chromatograms of the samples were aligned and deconvoluted using the open-source software MS-DIAL 4.9. The data were normalized using the MS-DIAL mTIC method, and the intensity of each compound was extracted and averaged. The peak annotation was performed based on the RI and spectral similarity matching, utilizing an in-house EI spectral library (Misra, 2019). The annotations were assigned at level 2 (putative annotation based on spectral library similarity) or level 3 (putatively characterized compound class based on spectral similarity to known compounds of a chemical class), following the classification proposed by Sumner et al. (2007).

b. Eco-evolutionary model of local adaptation

Changes in plant height H of each population i over time t were modelled as

$$\frac{dH_{i}}{dt}=H_{i}\times\mu_{i}\times(\rho_{i}-\varepsilon)$$

considering plant growth and plant response to climate. Stochasticity $\varepsilon$ was introduced from a normal distribution with mean 2.5 10^-3^ and sd 1 10^-2^. Plant growth rate $\mu$ was derived from marginal means contrasts between years at each site i. Climate sensitivity $\rho$ of each population i was calculated as

$\rho_{i}=\frac{\eta_{s}\left( \eta_{2020}-\eta_{1920} \right)}{100}+\frac{\theta_{s}\left( \theta_{2020}-\theta_{1920} \right)}{100}$,

where sensitivity to precipitation $\eta_{s}$ and sensitivity to temperature $\theta_{s}$ were derived from Principal Component Analysis (see also R code) and weighted for differences in precipitation and temperature between two sampling years (i.e., 2020 and 1920) at each site i. Initial conditions were plant average height as reported in Clausen et al. (1948). The model was run for 100 years from 1920 to 2020 with one year interval using automatic switching solver for Ordinary Differential Equations (ODE) (Soetaert et al., 2010).

**Supplementary Tables**

**Table S1** Description of study sites reporting occurrence (O), elevation ([m a.s.l.]), temperature (T [celcius]) and precipitation (P [mm/year]) in 1920 and 2020, and the relative differences (Δ) between the two years.

| **Site** | **O** | **Elevation** | **T 1920** | **T 2020** | **Δ T [%]** | **P 1920** | **P 2020** | **Δ P [%]** |
| --- | --- | --- | --- | --- | --- | --- | --- | --- |
| Bodega Bay | 1 | 8 | 12.06 | 13.44 | + 11.44 | 788.42 | 852.93 | + 8.18 |
| San Gregorio | 1 | 50 | 13.11 | 14.11 | + 7.63 | 645.92 | 731.01 | + 13.17 |
| Clayton | 1 | 201 | 16.11 | 16.89 | + 4.84 | 463.04 | 570.23 | + 23.15 |
| Knights Ferry | 0 | 90 | 16.83 | 17.39 | + 6.14 | 363.73 | 345.95 | – 5.03 |
| Groveland | 1 | 915 | 13.17 | 13.78 | + 4.63 | 1766.06 | 1330.71 | – 24.65 |
| Mather | 1 | 1400 | 11.17 | 11.83 | + 5.91 | 660.65 | 760.98 | + 15.19 |
| Aspen Valley | 1 | 1950 | 5.89 | 7.00 | + 18.85 | 831.34 | 783.59 | – 5.75 |
| Yosemite Creek | 1 | 2200 | 9.11 | 9.83 | + 7.90 | 1148.84 | 1227.07 | + 6.81 |
| Tuolumne Meadow | 1 | 2620 | 6.00 | 7.11 | + 18.50 | 1104.65 | 1040.38 | – 5.82 |
| Spuller Lake | 1 | 3150 | 5.11 | 6.22 | + 23.82 | 1094.49 | 1178.05 | + 7.63 |

**Table S2** Yarrow data (mean plant height and, standard error, and number of plants measured) reported in Clausen *et al.* (1948).

| **Site** | **Height** | **SE** | ***n.* plants** |
| --- | --- | --- | --- |
| BodegaBay | 45.3 | 1.23 | 59 |
| SanGregorio | 95.8 | 2.12 | 59 |
| Clayton | 63.4 | 1.42 | 57 |
| KnightsFerry | 67.4 | 1.92 | 57 |
| Groveland | 83.4 | 2.19 | 32 |
| Mather | 75.6 | 1.81 | 58 |
| AspenValley | 49.3 | 1.68 | 52 |
| YosemiteCreek | 49.1 | 1.66 | 53 |
| TenayaLake | 32.2 | 1.18 | 57 |
| TuolumneMdw | 20.8 | 1.11 | 46 |
| SpullerLake | 15.3 | 1.47 | 9 |
